# Application of Digital Colorimeter for Preliminary Characterization of Gold Nanoparticle Swarms Produced by *Termitomyces heimii* Using a novel Bioinspired Microfluidics Assay

**DOI:** 10.1101/2020.07.15.204453

**Authors:** Sujata Dabolkar, Nandkumar M. Kamat

## Abstract

In our laboratory work extending over several years we have successfully studied the biogeochemical role of termite mounds and their occupants the termites and the exosymbiont fungus-*Termitomyces*. Fungi appear to be promising for large scale production of nanoparticles (NPs) as these are simpler to grow both in laboratory and at industrial scale. This paper reports a novel microfluidic based assay system to detect Gold bioreduction capacity of different tissues in tissue based and cell free environment. Using sterile microtest wells, different tissues such as umbo, pileus, lamellae, stipe context, stipe epicutis, pseudorrhiza context, pseudorrhiza epicutis of *Termitomyces heimii* mature fruitbodies were tested with 200μl chloroauric acid (one mM) and after an interval of 5, 10, 15, 30, 45, 60, 120 min and 12, 24 and 48 hours. The results in terms production of distinct nanoparticles were directly visualized microscopically and using mobile based digital colorimeter. Membrane filtered sterile water soluble extracts (SWSE) from the same tissues were similarly screened. The results manifested by mono and polydisperse GNPs and microparticles of mixed size groups demonstrated that cell free system can be potentially useful for bioinspired fabrication of GNPs. Further work in this direction is in progress using several *termitomyces* pure cultures.

## Introduction

Poor and under developed countries like India and resource starved universities find it difficult to have easy access to expensive instrumentation for characterization of Gold nanoparticles (GNPs). However now the new Apps such as digital colorimeter from laboratory tools make it possible to rapidly detect GNP formation and perform quick and simple analysis. Having seen the chemical creativity of termitophilic mushrooms in our laboratory we aimed to use microfluidic assay for rapid screening of gold bioreduction system in this species. *Termitomyces heimii* being the most dominant species and state mushroom of Goa we used this for our study. However, it was not easy to rapidly detect and characterize swarms of gold micro and nanoparticles. It was at this point that we came across a mobile based digital colorimeter which was found useful in analysis of the GNP swarms and obtain spectra in visible band. Swarm are formed by the collective behavior of GNPs. Inspired by animal interactions, the autonomous movement and collective behavior of synthetic nanomaterials are of considerable interest as they have implications for the future in nanomachinery, nanomedicine, and chemical sensing (Kagan et al., 2011). Values such as CIE LAB, Chroma, Hue°, RGB, color names, real time visible spectra (400 nm to 700 nm) can be recorded using this digital tool. The CIE LAB color space (also known as CIE L*a*b* or sometimes abbreviated as simply “Lab” color space) is a color space defined by the International Commission on Illumination (CIE) in 1976. It expresses color as three values where L* stands for the lightness from black to white, a* from green to red, and b* from blue to yellow. Chroma (Saturation) may be defined as the strength or dominance of the hue, the quality of a color’s purity, intensity or saturation (Solomon & Breckon 2011). Hue is common distinction between colors positioned around a color wheel. On the outer edge of the hue wheel are the intensely saturated hues whereas towards the center of the color wheel no hue dominates and becomes less and less saturated. The RGB color model is an additive color model in which red, green and blue lights are added in various ways to reproduce a broad array of colors (Meruga et al., 2014). Finally, it has been shown that absorbance peaks of GNPs are correlated to their size and we aimed to test the ability of digital colorimeter to get an idea of size distribution of GNPs in the swarms.

Statistical importance of the data was obtained using jvenn, a new JavaScript library (http://bioinfo.genotoul.fr/jvenn/example.html) which processes lists and produces Venn diagrams. Venn diagrams with more than four lists, are much harder to interpret. To solve this problem, the classical or Edwards-Venn representation introduces new shapes providing a clearer view (Philippe Bardou et al., 2014). Jvenn enables to compare up to six lists and updates the diagram automatically when modifying the lists content.

## Methodology

### Sample collection

*Termitomyces heimii* being the most dominant species in Goa and state mushroom of Goa we use this for our study. Fresh, healthy specimens of *Termitomyces heimii* Natarajan (1979) were collected from fields of Taleigao, Goa during monsoon season, 2019 and taxonomically identified using standard published *Termitomyces* keys (Heim R. 1942,1977; Natarajan,1979) (**Fig.1**). Dried herbarium is deposited in Goa university mycological herbarium collection.

**Fig.1.**
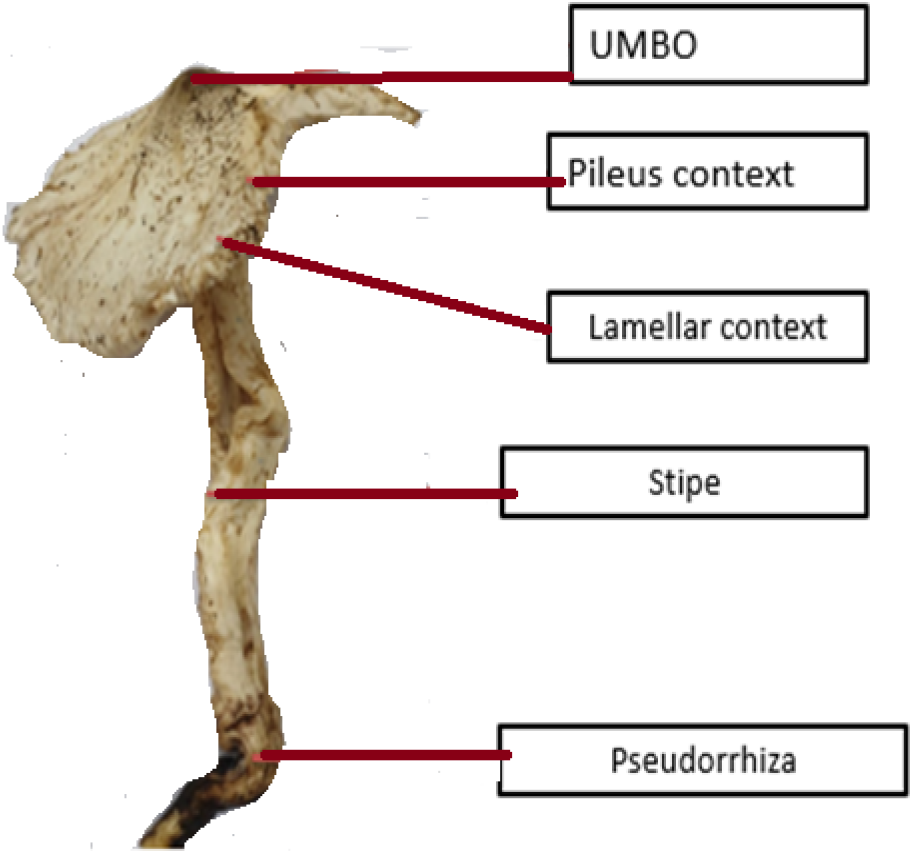
Scheme for specific processing of different fruitbody parts

### Processing of the specimens

*T. heimii* specimens were cleaned with 95% ethanol (v/v) upto 30 seconds and photographed. Specific processing of each part of the fruitbody ie umbonal tissue, pileus context, lamellae, stipe context, stipe epicutis, pseudorrhiza context, pseudorrhiza epicutis was carried out. Using sterile forceps small pieces of the tissues were transferred into a microtest plate (Tarson, Mumbai) with 96 wells having volumetric capacity of 420 μl under a laminar air flow bench. Care was taken to use identical tissue fragments appropriate equivalent to 200 μm size. Tissues were tested with 200 μl Chloroauric acid (one mM) (**Fig. 2**) and after an interval of 5, 10, 15, 30, 45, 60, 120 minutes and 12, 24 and 48 hours. Nine replicates of each tissue were used.

**Fig.2.**
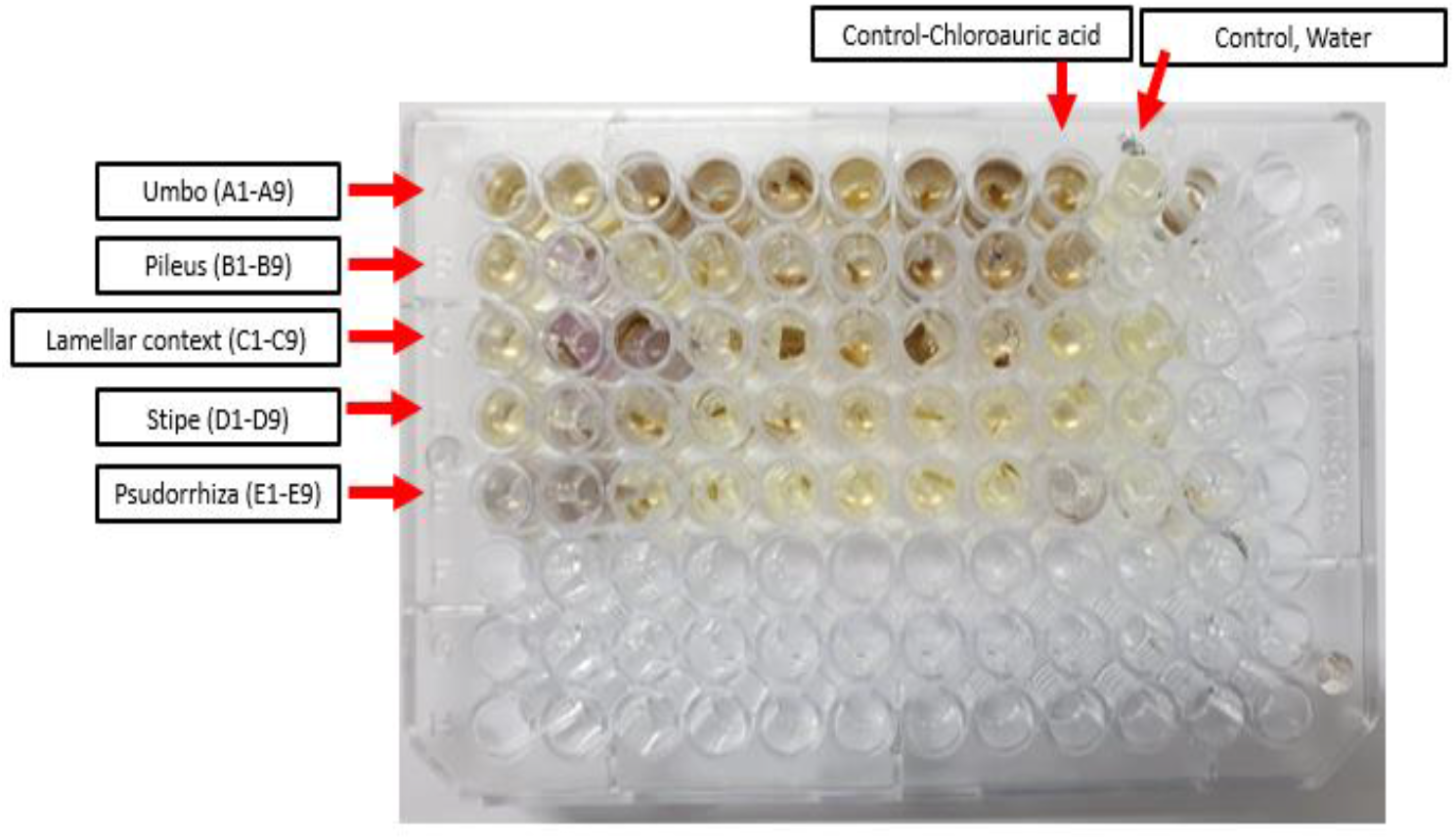
Design of Microfluidics based assay using Microtest plate

### Preparation of SWSE

Sterile water soluble extracts (SWSE) (**Fig. 3**) were prepared by grinding in sterile mortar with pestle, centrifuged and membrane filtered (0.22 μm pore size, 30 mm diameter-HIMedia laboratories). The SWSE were stored at refrigerated temperature in sterile test tubes. The extracts (210 μl) and chloroauric acid (210 μl) were mixed in equal proportion in the wells of microwell test plate and checked after interval of 5, 10, 15, 25, 30, 45, 60, 120 minutes and after 12, 24, 48 hours. The assay design was similar to fig.3.

**Fig.3.**
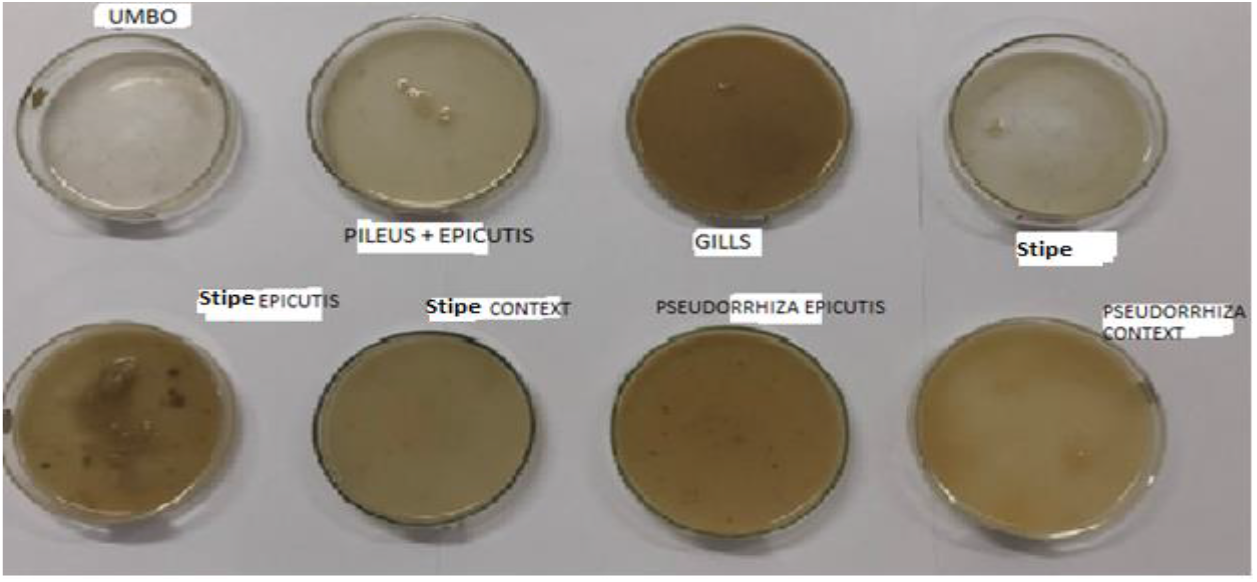
Homogenized aquous extracts from different *T. heimii* fruit body tissue

### Stereomicroscopic visualization of swarms

The microtest plate with the GNP swarms was visualized under stereomicroscope (Olympus SZ51, model SZ2-ILST, olympus corporation, Tokyo, Japan) (**Fig.4**). Care was taken to bring the swarm view under uniform illumination in bright light.

**Fig.4.**
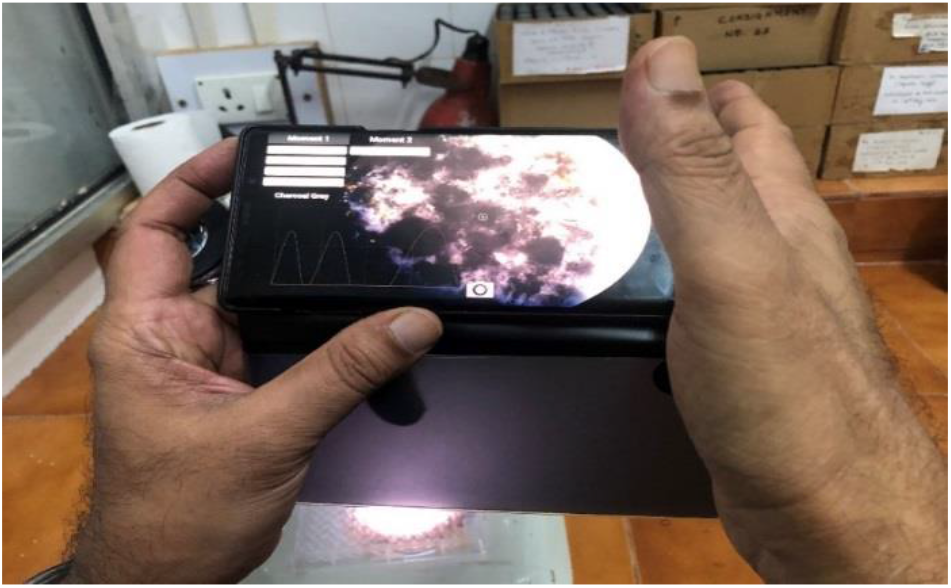
Direct Acquisition of images for stereomicroscopic characterization

### Use of Digital Colorimeter App

Scanning of the swarms was done with 12 MP plus dual rear (F 1.5/ F 2.4) camera on Samsung Galaxy Note 9 with colorimeter software (http://researchlabtools.blogspot.com/) (Ravindranath et al, 2018) version 3.5.2, developed by Research Lab Tools, São Paulo, Brazil.

### Digital color analysis and colorimetric data

The color terminology is used according to color data based of the App. Colorimeter software was used to record values such as CIE LAB, Chroma, Hue°, RGB, color names, real time visible spectra (400 nm to 700 nm). The App allows online and offline analysis of samples.

### Use of Venn diagrams

Venn diagrams are commonly used to display list comparison. However, when the number of input lists exceeds four, the diagram becomes difficult to read. Alternative layouts and dynamic display features can improve its use and its readability. The jvenn library accepts three different input formats “Lists”, “Intersection counts” and “Count lists”. For “Intersection counts”, the lists are given a label (“A” or “B”) which is used to make the correspondence between the list and its count. Finally, “Count lists” provide a count number for each element of a list. Hence, with “Count lists” the figures presented in the diagram correspond to the sums of counts of all elements shared between lists For “Lists” and “Count lists”, jvenn computes the intersection counts and displays the chart (http://bioinformatics.psb.ugent.be/). Vein diagrams were plotted using tissues and lambda max values.

## Results

Fresh, healthy *Termitomyces heimii* which is dominant species in Goa were successfully obtained and were taxonomically identified using standard published *Termitomyces* keys (Heim R. 1942, 1977; Natarajan,1979; DeSouza & Kamat, 2018, 2019). GNP swarms were detected in all treatments and could be visualized easily under stereomicroscope (**Fig.5 & Fig 6**). Umbonal tissue produced grey GNP swarms and color values as shown in table 1. Overall the color range is from grey to juniper green. The chromaticity values showed difference and chroma values ranged from 4 to 47 whereas Hue differed from 40 to 199. The R value varied from 131 to 177, G from 102 to 187 whereas B from 27 to 191. Detail treatment of absorbance value is given in table 3. Similarly the colors and color analysis and absorbance values of other tissues and SWSE are shown in **Table 1** and **Table 2**.

**Table 1:**
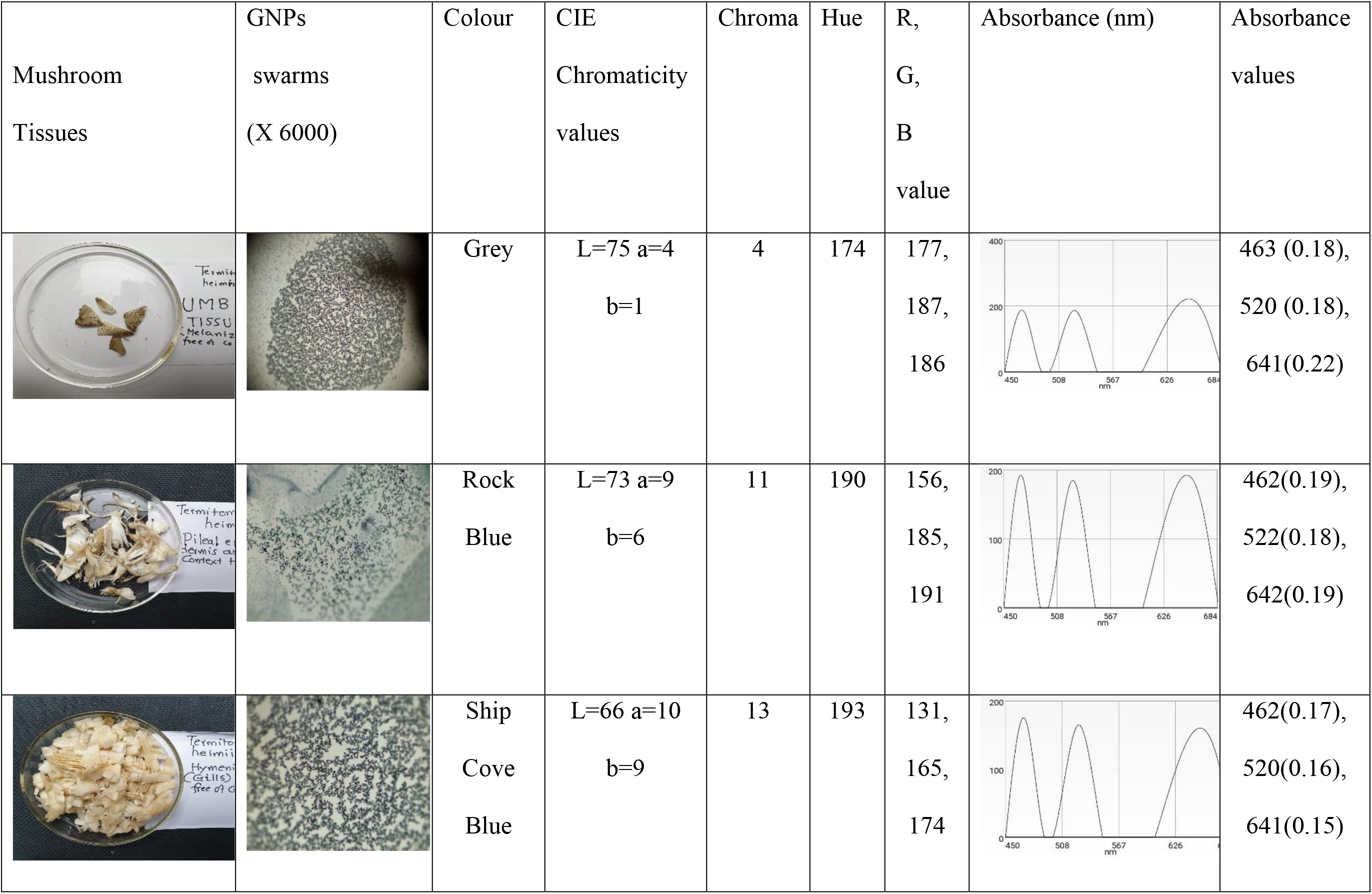
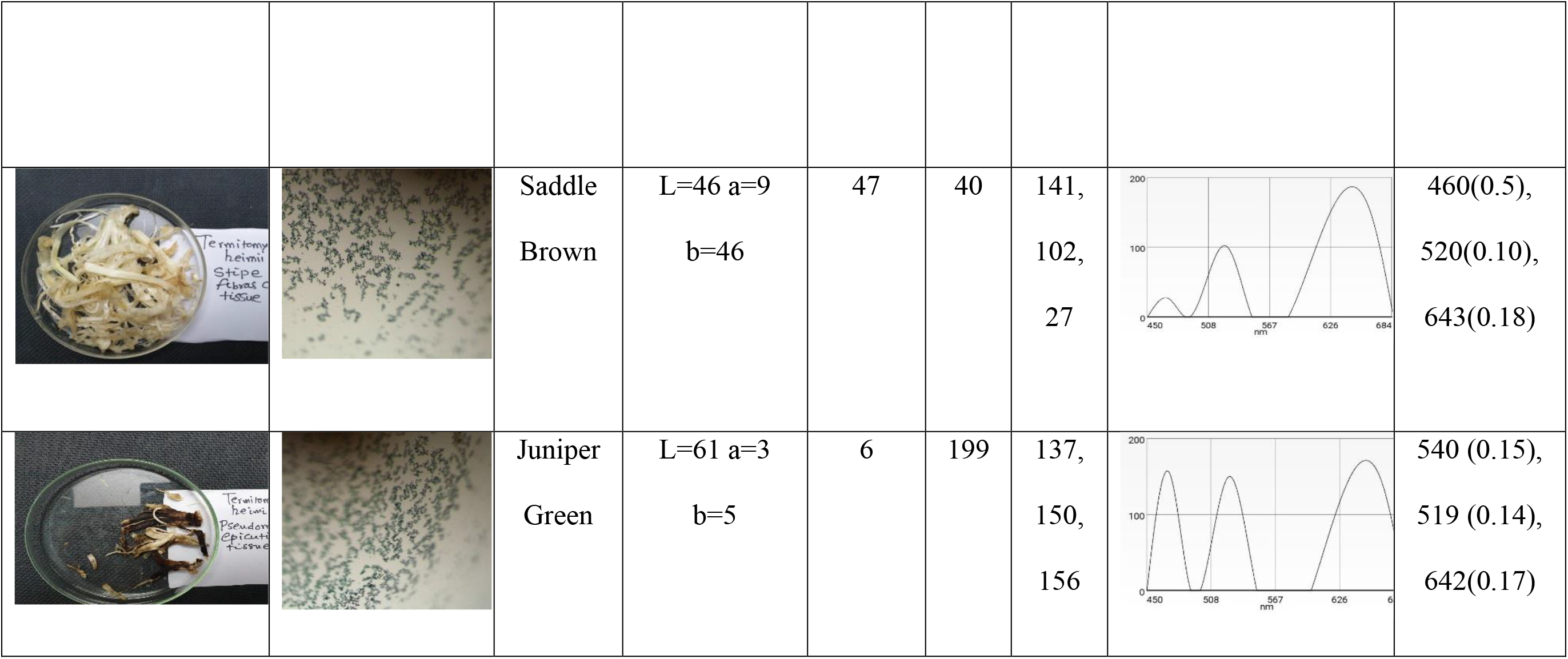
Colour analysis and Colorimetric absorption characterization of presumptive GNP swarms

| Mushroom<br>Tissues | GNPs<br>swarms<br>(X 6000) | Colour | CIE<br>Chromaticity<br>values | Chroma | Hue | R,<br>G,<br>B<br>value | Absorbance (nm) | Absorbance<br>values |
| --- | --- | --- | --- | --- | --- | --- | --- | --- |
|  |  | Grey | L=75 a=4<br>b=1 | 4 | 174 | 177,<br>187,<br>186 |  | 463 (0.18),<br>520 (0.18),<br>641(0.22) |
|  |  | Rock<br>Blue | L=73 a=9<br>b=6 | 11 | 190 | 156,<br>185,<br>191 |  | 462(0.19),<br>522(0.18),<br>642(0.19) |
|  |  | Ship<br>Cove<br>Blue | L=66 a=10<br>b=9 | 13 | 193 | 131,<br>165,<br>174 |  | 462(0.17),<br>520(0.16),<br>641(0.15) |
|  |  | Saddle<br>Brown | L=46 a=9<br>b=46 | 47 | 40 | 141,<br>102,<br>27 |  | 460(0.5),<br>520(0.10),<br>643(0.18) |
|  |  | Juniper<br>Green | L=61 a=3<br>b=5 | 6 | 199 | 137,<br>150,<br>156 |  | 540 (0.15),<br>519 (0.14),<br>642(0.17) |

**Table 2:**
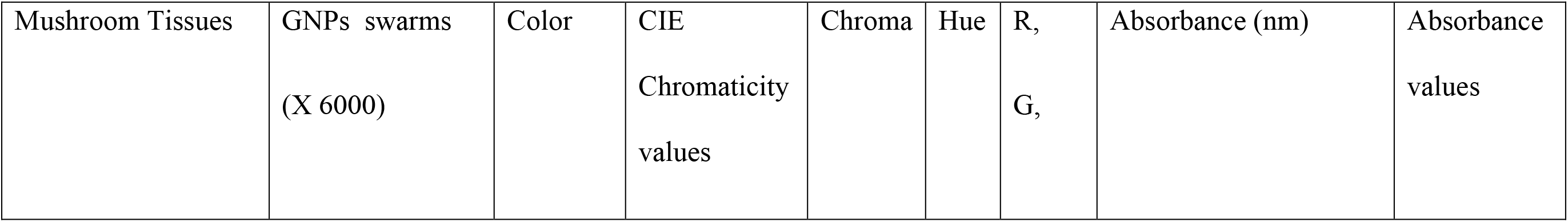
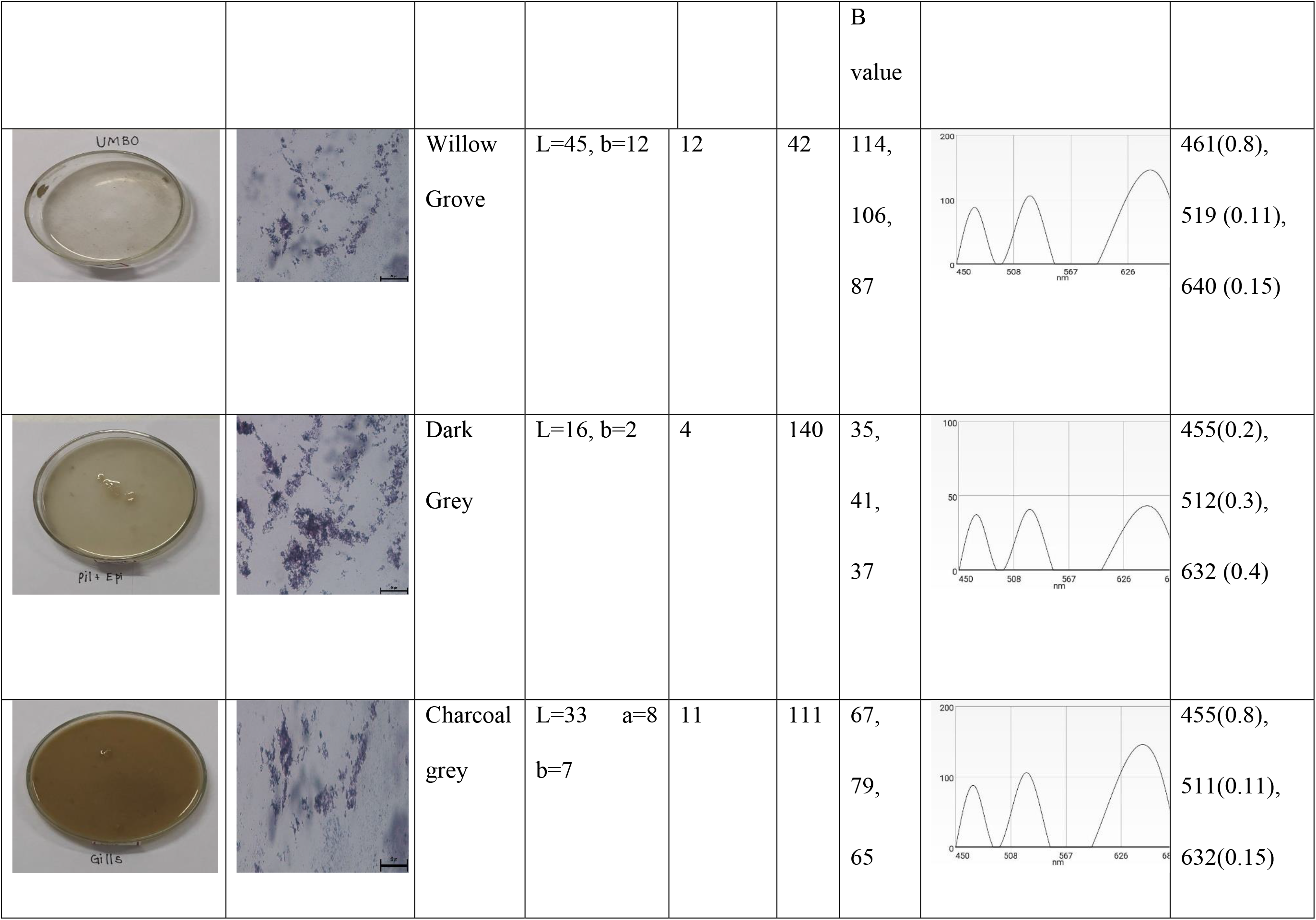
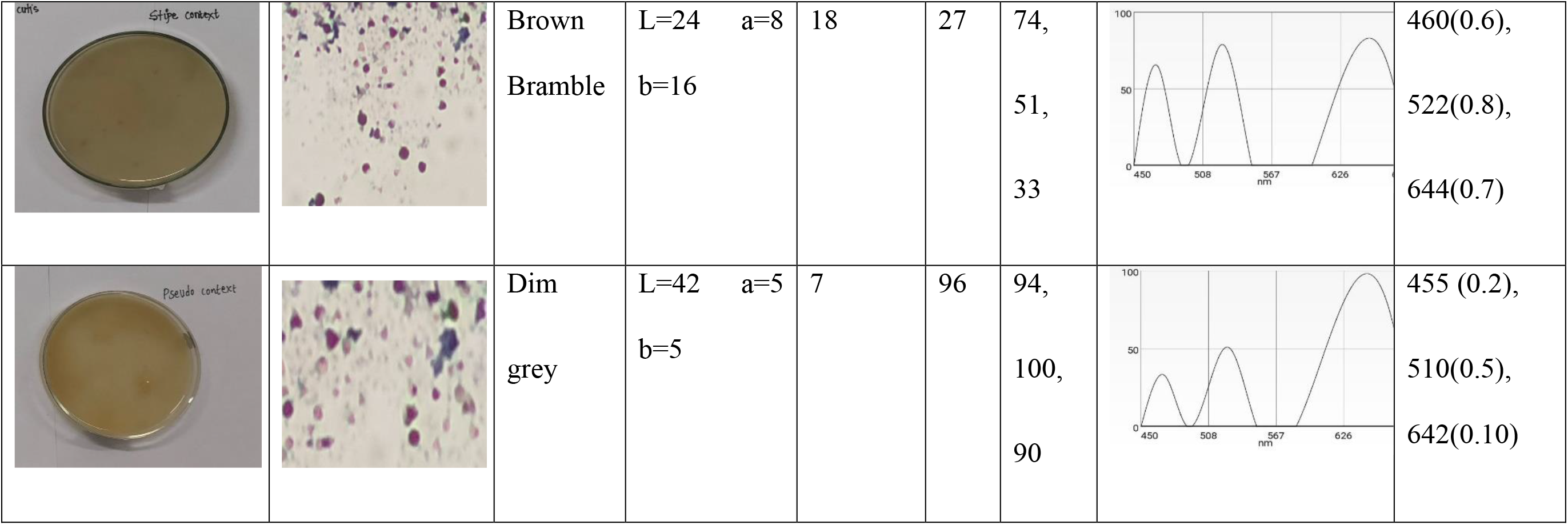
Colour analysis and Colorimetric absorption characterization of presumptive GNP swarms

| Mushroom Tissues | GNPs swarms<br>(X 6000) | Color | CIE<br>Chromaticity<br>values | Chroma | Hue | R,<br>G, | Absorbance (nm) | Absorbance<br>values |
| --- | --- | --- | --- | --- | --- | --- | --- | --- |

|  |  |  |  |  |  | B<br>value |  |  |
| --- | --- | --- | --- | --- | --- | --- | --- | --- |
|  |  | Willow<br>Grove | L=45, b=12 | 12 | 42 | 114,<br>106,<br>87 |  | 461(0.8),<br>519 (0.11),<br>640 (0.15) |
|  |  | Dark<br>Grey | L=16, b=2 | 4 | 140 | 35,<br>41,<br>37 |  | 455(0.2),<br>512(0.3),<br>632 (0.4) |
|  |  | Charcoal<br>grey | L=33 a=8<br>b=7 | 11 | 111 | 67,<br>79,<br>65 |  | 455(0.8),<br>511(0.11),<br>632(0.15) |
|  |  | Brown<br>Bramble | L=24<br>b=16<br>a=8 | 18 | 27 | 74,<br>51,<br>33 |  | 460(0.6),<br>522(0.8),<br>644(0.7) |
|  |  | Dim<br>grey | L=42<br>b=5<br>a=5 | 7 | 96 | 94,<br>100,<br>90 |  | 455 (0.2),<br>510(0.5),<br>642(0.10) |

**Table 3:** Visible spectral characteristics of GNP swarms using *T. heimii* tissue sample and SWSE

| Absorbance<br>wavelength<br>(nm) | Tissue |  |  |  |  | SWSE |  |  |  |  |
| --- | --- | --- | --- | --- | --- | --- | --- | --- | --- | --- |
|  | I<br>Umbonal<br>tissue | II<br>Pileus<br>context | III<br>Lamellae | IV<br>Stipe<br>context | V<br>Pseudorrhiza<br>context | VI<br>Umbonal<br>context | VII<br>Pileus<br>context | VIII<br>Lamellae | IX<br>Stipe<br>context | X<br>Pseudorrhiza<br>context |
| 455 | - | - | - | - | - | - | + | + | - | + |
| 460 | - | - | - | + | - | - | - | - | + | - |
| 461 | + | - | - | - | - | - | - | - | - | - |
| 462 | + | + | - | - | - | - | - | - | - | - |
| 463 | + | - | - | - | - | - | - | - | - | - |
| 510 | - | - | - | - | - | - | - | - | - | + |
| 511 | - | - | - | - | - | - | - | + | - | - |
| 512 | - | - | - | - | - | - | + | - | - | - |
| 519 | - | - | - | - | + | + | - | - | - | - |
| 520 | + | - | + | + | - | - | - | - | - | - |
| 522 | - | + | - | - | - | - | - | - | + | - |
| 540 | - | - | - | - | + | - | - | - | - | - |
| 632 | - | - | - | - | - | - | + | + | - | - |
| 640 | - | - | - | - | - | + | - | - | - | - |
| 641 | + | - | + | - | - | - | - | - | - | - |
| 642 | - | + | - | - | + | - | - | - | - | + |
| 643 | - | - | - | + | - | - | - | - | - | - |
| 644 | - | - | - | - | - | - | - | - | + | - |

**Fig.5.**
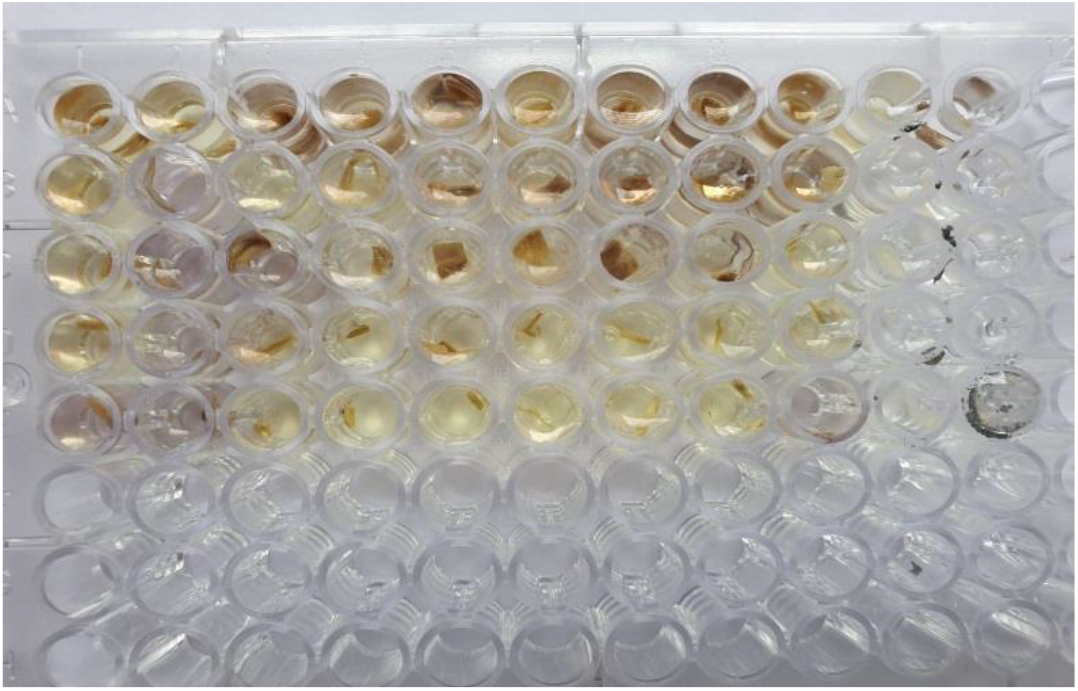
Positive bioreduction obtained with homogenized tissues as indicated by color changes

**Fig.6.**
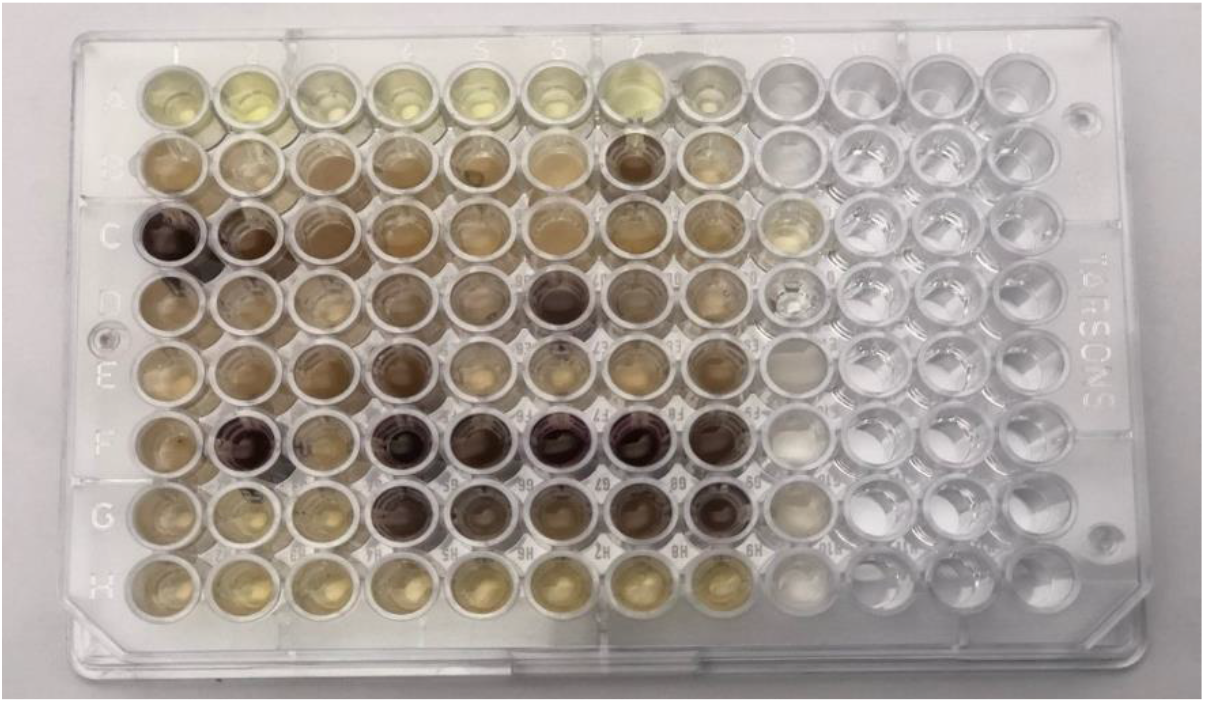
Positive bioreduction with cell free membrane filtered SWSE indicated by change in colour, yellow is control

**Table 3** shows 30 different peaks obtained using each tissue and extract. Absorbance value ranged from 455 nm to 644 nm. Stipe exhibited most promising results with peaks at 455, 510, 642 nm. **Table 4** gives approximate GNP size range diameter which ranged from 5 nm to 100 nm. It was noticed that only extract system was producing GNP swarms at wavelength of 455 nm, whereas only umbonal tissue produced maximum absorption at 461 nm similar results were obtained in remaining reaction as shown in table 3. The correlation between the absorbance values verses the tissue based system and cell free environment is shown (**Fig7a-7f**).

**Table 4:** Approximate size range of GNP swarms produced using *T. Heimii*

| **Wavelength reported in present assay (nm)** | **Approximate GNP size range (nm)** | **Reference** |
| --- | --- | --- |
| 455, 460, 461, 462, 463 | < 5 | Ted Pella Inc. |
| 510, 511 | 5 | Ted Pella Inc. |
| 512, 519 | 10 | Sigma-Aldrich |
| 520, 522 | 15 | Sigma-Aldrich, Ted Pella Inc. |
| 540 | 10-30 | Radtsig et al., 2016 |
| 632 | 50-80 | Ted Pella Inc. |
| 640 | 80-100 | Sigma-Aldrich |
| 641 | 100 | Sigma-Aldrich |
| 642, 643, 644 | 100 | Sigma-Aldrich, Ted Pella Inc. |

**Fig.7 (a-f):**
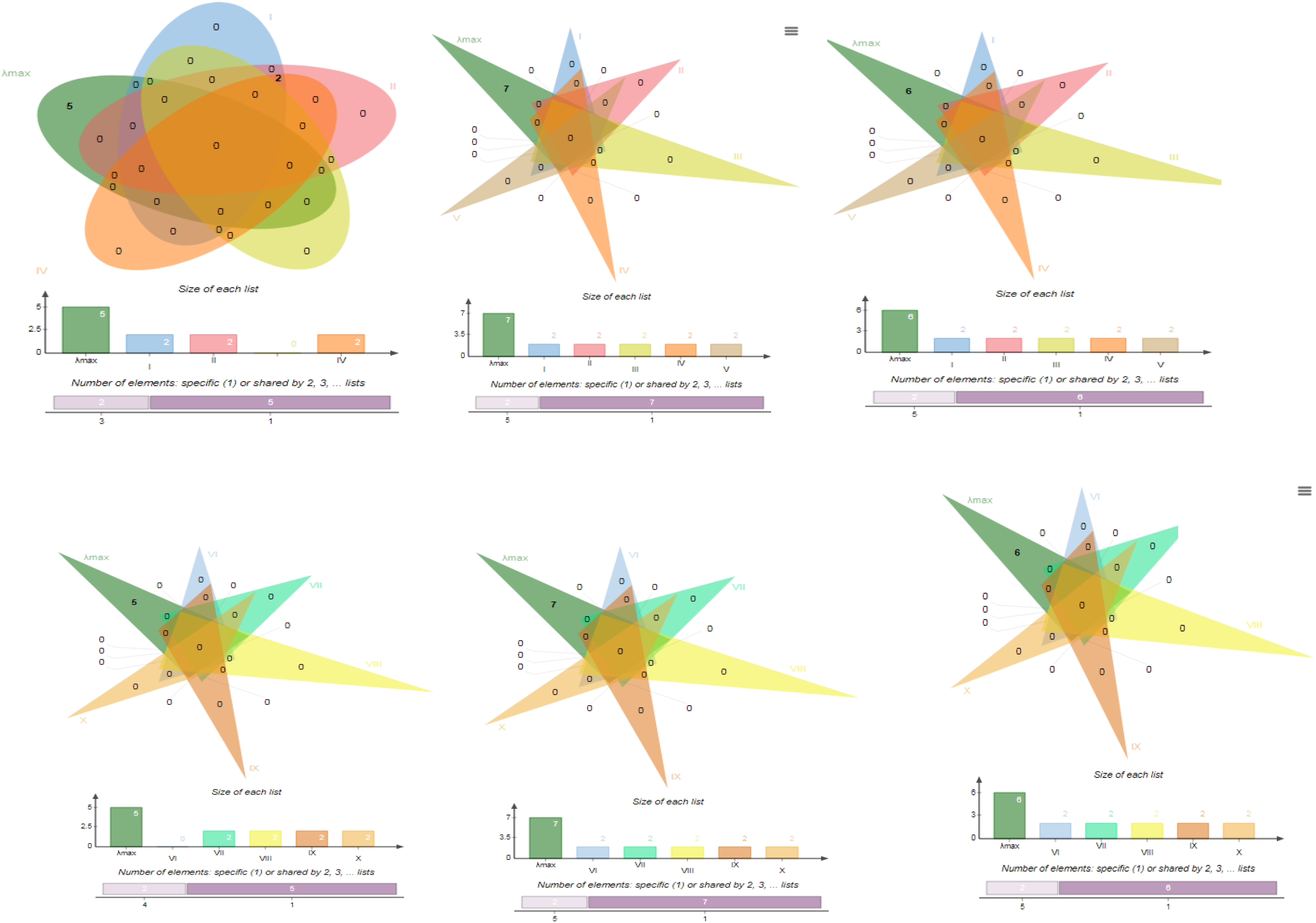
7(a-c)- The shape corresponding to the lists involved in the intersection are highlighted in case of tissue, (a-455-463 λmax, b-510-540,c-632-644). 7(d-f)- The shape corresponding to the lists involved in the intersection are highlighted in case of extracts (a- 455-463 λmax,b-510-540,c-632-644).

## Discussion

This paper reports a novel microfluidics based assay system to detect Gold bioreduction capacity of different tissues in Termitophilic mushrooms (Kalia & Kaur, 2018; DeSouza & Kamat, 2017; de Souza & Kamat, 2018, 2019) in tissue based and cell free environment. Umbonal tissue, pileus context, lamellae, stipe context, stipe epicutis, pseudorrhiza context, pseudorrhiza epicutis of *Termitomyces heimii* mature fruitbodies successfully produced GNPs for the first time. We were successful in producing membrane filtered SWSE from same tissues and also successful in producing GNPs from the same extracts. Our assay can be useful to carry out large number of replicates, under sterile forms. Microtest plates can be directly visualized due to its transparent makeup and swarms can be directly characterized under stereomicroscope. Small amounts of gold solutions and small amount of SWSE can be tested this microfluidic assay. Using rapid screening of large number of biological or microbiological gold bioreduction systems.

In case of tissues the colour varied from grey, rock blue, ship cove blue, juniper green to saddle brown where as in case of SWSE it was willow grove, drim grey dark grey, charcoal grey to brown gramble. For small (~30 nm) monodisperse gold nanoparticles, the surface plasmon resonance phenomenon causes an absorption of light in the blue-green portion of the spectrum (~450 nm) while red light (~700 nm) is reflected, yielding a rich red color. As particle size increases, the wavelength of surface plasmon resonance related absorption shifts to longer, redder wavelengths (https://www.sigmaaldrich.com/). Larger the size, darker is the color and may also shift to blue in case of colloidal particles (Jana et al., 2001; Haiss et al., 2007; Martinez et al., 2012). Absorbance reading tells the composition and size of NPs (Doak et al., 2010). SWSE prepared from same tissue do not produce GNPs of same size or with same concentration. The GNP swarm population is represented by 18 different size groups ranging from less than 5 nm to 100 nm. The concentration of nanoparticles as function of optical absorbance ranges from 0.12 to 0.8 indicating that some bioreduction systems are much more efficient in production of GNPs this includes GNPs of the size of less than 5 to 15 nm.

It was found that stipe was showing most promising results in case of both tissue and SWSE. CIE chromaticity values ranged from L (16-75), a (3-33) and b (1-46) thus indicating the lightness from black to white, green to red and blue to yellow (Cheng et al.,2014). Choma values ranged from 4 to 47 and hue values from 40 to 199 indicating the strength or dominance of the hue, the quality of a color’s purity, intensity or saturation.

GNPs produced using tissue showed low absorbance from min 0.10 to max 0.5 whereas SWSE produced GNP with absorbance ranging from 0.1 to 0.8 thereby indicating a more efficient system in cell free environment. Low absorbance 0.19 for intact tissue indicating low concentration of GNP swarms. Mean absorbance produced by treatment with extract 0.40 indicating almost double the bioreduction efficiency of intact tissue in case of GNP production. Thus, cell free environment is much better system to produce polydisperse GNP swarms in higher concentration.

In all 30 different peaks ranging from 455 to to 644 nm were obtained using each tissue and extract thus indicating the presence of nanoparticle of size 5 nm to 100 nm (https://www.sigmaaldrich.com). Mushrooms are rich in proteins and have high availability of the amino acids lysine, tryptophan, glutamic acid and aspartic acid (Hsu et al., 2002). It is also reported that certain mushroom extract contain polysaccharide/oligosaccharide complex (Cho et al., 2003). FTIR studies have also shown the possible biomolecules responsible for capping and efficient stabilization of the metal nanoparticles synthesized using mushroom extract (Philip, 2009). It was noticed certain SWSE and tissues produced specific wavelength for example umbonal, pileaus context, pseudorrhizal context extract system was producing GNP swarms at wavelength of 455 nm and only umbonal tissue produced at 461 nm. Similar results were obtained in remaining reaction as shown in table 3. It was noticed that there is relationship between solubility and swarm formation and it could be a different molecule based bioreduction system.

Preliminary characterization of GNP swarms is important as in high throughput screening system one cannot differentiate the most promising system and it can be time consuming. Once you carry out the preliminary results you zero down to the specific system to obtain the promising system and then can go for final characterization of the GNPs. Preliminary results can also help in standardization of the procedure and also there is no waste of resources. In poor and under developed countries like India and resource starved universities researchers find it difficult to have easy excess to expensive instrumentation for characterization of GNPs. However now the new Apps such as digital colorimeter from developers Laboratory tools make it possible to rapidly detect GNP formation and perform quick analysis.

## Conclusions

Our work clearly demonstrates that simple and easy to use mobile digital Colorimeter Apps can be used for primary optical characterization of swarms of Gold nanoparticles. This is useful in rapid screening of large number of microbiological gold bioreduction systems. The spectral absorbance profile detected in visible range also helps in understanding the presumptive size of GNPs in swarms. For high throughput screening systems we recommend development of more such mobile based apps. Our present approach has helped us to fabricate a very sensitive Gold biosensor. Pure mycelial cultures of *T. heimii* also produced identical results. This would be published separately.

## Acknowledgements

The authors would like to thank RNSB project for the support. This work was also supported by UGC SAP Phase III Biodiversity, Bioprospecting programme and Goa University Fungus Culture Collection GUFCC). First author also acknowledges UGC, NF OBC Junior Research fellowship. Thanks for guidance from Dr. Absar Ahmad director, interdisciplinary center for Nanotechnology AMU regarding potential of GNPs.

## Figures and Tables

**Fig.1.** Scheme for specific processing of different fruitbody parts

**Fig.2.** Design of Microfluidics based assay using Microtest plate

**Fig.3**. Homogenized aqueous extracts from different tissues

**Fig.4**. Direct Acquisition of images for stereomicroscopic characterization

**Fig.5.** Positive bioreduction obtained with homogenized tissues as indicated by color changes

**Fig.6**. Positive bioreduction with cell free membrane filtered SWSE indicated by change in colour, yellow is control

**Fig.7 (a-f)**. 7(a-c): The shape corresponding to the lists involved in the intersection are highlighted in case of tissue, (a-455-463, b-510-540, c-632-644 λmax). 7(d-f): The shape corresponding to the lists involved in the intersection are highlighted in case of extracts, (a-455-463 λmax, b-510-540, c-632-644 λmax).

